# Emotional ego- and altercentric biases in high-functioning autism spectrum disorder: Behavioral and neurophysiological evidence

**DOI:** 10.1101/2021.11.12.468198

**Authors:** Helena Hartmann, Lukas Lengersdorff, Hannah H. Hitz, Philipp Stepnicka, Giorgia Silani

**Author notes:** Corresponding authors, Department of Clinical and Health Psychology, University of Vienna, Liebiggasse 5, 1010 Vienna, Austria, Social, Cognitive and Affective Neuroscience Unit, Department of Cognition, Emotion, and Methods in Psychology, University of Vienna, Liebiggasse 5, 1010 Vienna, Austria. Shared first authorship (Helena Hartmann, Lukas Lengersdorff).

## Abstract

Self-other distinction is a crucial aspect of social cognition, as it allows us to differentiate our own mental and emotional states from those of others. Research suggests that this ability might be impaired in individuals with autism spectrum disorder (ASD), but convincing evidence of self-other distinction deficits in the emotional domain is lacking. Here we aimed at evaluating emotional self-other distinction abilities in adults with and without ASD, in two behavioral pilot studies and one fMRI study. By using a newly developed virtual ball-tossing game that induced simultaneous positive and negative emotional states in each participant and another person, we were able to measure emotional egocentric and altercentric biases (namely the tendency to ascribe self-/other-related emotions to others/ourselves, respectively). Despite no behavioral differences, individuals with ASD showed decreased activation 1) in the right temporoparietal junction (rTPJ) during active overcoming of the emotional egocentric bias vs. passive game viewing, and 2) in the right supramarginal gyrus (rSMG) during ego-vs. altercentric biases, compared to neurotypical participants. These results suggest a different recruitment of these two regions in ASD when dealing with conflicting emotional states of oneself and another person. Furthermore, they highlight the importance of considering different control conditions when interpreting the involvement of rTPJ and rSMG during self-other distinction processes.

## 1 Introduction

Social cognition, the capacity to sense, represent and judge our own social behaviors and those of others, is an ubiquitous aspect of the human mind and crucial for everyday social interactions (see Frith & Frith, 2008 for a review). Humans represent and infer other’s mental states – an ability known as mentalizing or Theory of Mind – using multiple self-related processes: e.g. when putting oneself in the shoes of another person (Baron-Cohen et al., 2013; Happé et al., 2017; Singer, 2006 for reviews). At the same time, adequate social behavior demands that we can distinguish between self- and other related representations, a crucial cognitive skill termed self-other distinction (de Vignemont & Singer, 2006; Preckel et al., 2018 for reviews; Reniers et al., 2014; Thirioux et al., 2014).

The importance of self-other distinction becomes evident in cases where it breaks down. For example, it has been proposed that this ability is disrupted or at least qualitatively different in individuals with autism spectrum disorder (ASD), a neurodevelopmental condition largely characterized by impairments regarding social communication and social interaction (American Psychiatric Association, 2013). ASD individuals, for example, are less likely to draw a sharp distinction between their own and another’s perspective (Bird & Viding, 2014 for a review), and are more likely to confuse self- and other-related emotions (de Guzman et al., 2015 for a review; Fan et al., 2014 for a meta-analysis; Gu et al., 2015), leading to distorted cognitive and affective representations named biases. Deschrijver & Palmer (2020) have recently suggested that insufficient monitoring of mental conflicts – i.e. when mental states differ between oneself and another person, might be the main reason behind the aforementioned difficulties in ASD, rather than the general lack of mental representation abilities or “mindblindness”. In spite of the abundance of studies on increased cognitive biases in ASD, using e.g. false belief or social discounting tasks (Begeer et al., 2012; Frith & de Vignemont, 2005 for a review; Nijhof et al., 2020; Tei et al., 2019), biases in the emotional domain seem to have gotten less attention over the years. However, affective social competences are often a major problem for individuals with ASD, and there is still a wide debate which components of social cognition are indeed impaired in this disorder (Deschrijver & Palmer, 2020 for a recent review; Dziobek et al., 2008; Mazza et al., 2014).

Behavioral and especially neural mechanisms underlying emotional self-other distinction and the occurrence of *egocentric* (i.e. the tendency to project one’s own states onto others) and *altercentric* (i.e. the tendency to absorb another’s state onto one’s own) biases in the emotional domain have only been investigated by few researchers up to this point, with single studies ranging from neurotypical (NT) children to older adults (Hoffmann et al., 2015; Hoffmann, Banzhaf, et al., 2016; Repacholi & Gopnik, 1997; Riva et al., 2016, 2019; Silani et al., 2013; Steinbeis et al., 2014; von Mohr et al., 2020). To date, only one study (Hoffmann, Koehne, et al., 2016) was performed to assess emotional self-other distinction in ASD. By using a visuo-tactile task where participants rated the (un)pleasantness of different objects simultaneously touching themselves and another participant, the authors observed preserved behavioral self-other distinction (but not mentalizing abilities) in ASD compared to matched NT individuals. Furthermore, in an independent pool of ASD individuals, they observed reduced resting state connectivity in the brain mentalizing network (i.e. right temporoparietal junction; rTPJ), but intact network connectivity of the right supramarginal gyrus (rSMG), a region which plays a crucial role in overcoming those biases (Hoffmann et al., 2015; Preckel et al., 2018; Silani et al., 2013). Despite being the first of its type, the study suffered of three shortcomings that need to be addressed: Firstly, the study did not clearly separate egocentric and altercentric biases, which may have led to them cancelling each other out, resulting in visibly preserved self-other distinction. Secondly, participants were tested in their pleasantness evaluation of simple touch stimuli, but the question remains whether the same holds true for more complex emotions evident in daily social interactions. Lastly, the study could not establish a direct link between behavior and brain as independent samples were used, making it difficult to draw conclusions involving both.

In order to replicate and extend this important initial work, we employed a newly developed version of the *Cyberball* task (a virtual ball-tossing game; Novembre et al., 2015; Williams & Jarvis, 2006), combined with functional Magnetic Resonance Imaging (fMRI) assessment. The task was designed to induce simultaneous positive and negative emotional states in a participant and another person by means of inclusion and exclusion from the game. Participants either took the role of active player or passive observer in the game, judged either their own emotional state or the one of the (spatially) opposite player and could either be in a congruent or incongruent emotional state (elicited through game inclusion or exclusion) with that other person. In short, the task entailed three different conditions: *self active* (= judging yourself as an active player), *other active* (= judging another person as an active player), and *other passive (=* judging another person but from a neutral, outside perspective as a passive observer). While the first two conditions were employed to resemble previous operationalizations of emotional egocentric biases (i.e. altercentric biases are subtracted from egocentric biases; Silani et al., 2013); the last condition was introduced to overcome the limitation of such operationalization (namely the impossibility to assess the egocentric and altercentric biases independently). By using an emotionally neutral condition, where participants are asked to judge other emotions without being themselves directly involved, we could control for task complexity (i.e., incongruency between different emotional states) without cancelling out the processing of self-other representations, and therefore capturing the emotional egocentric bias in a cleaner way.

The new task was first validated in two behavioral studies with independent NT samples (*n*’s = 45 and 52, see Supplementary Material) and then applied in our fMRI study, testing 21 NT and 21 ASD individuals. We hypothesized higher emotional egocentric biases (EEB) and/or emotional altercentric biases (EAB) in individuals with ASD compared to NTs, but no group differences during passive observation. In other words, we predicted lower self-other distinction abilities in the ASD compared to NTs, indicated by stronger ego- and altercentric judgment shifts when dealing with concurrent incongruent compared to congruent emotional states. As previous studies had used the EAB as a control, here we were additionally interested in differences between EEB and EAB. On the neurophysiological level, we hypothesized reduced activity in areas related to self-other distinction, such as rTPJ and rSMG in individuals with ASD compared to NTs during two active playing conditions, but no group differences in the passive condition.

## 2 Methods

### 2.1 Open data and materials statement

Unthresholded statistical maps of the fMRI data are uploaded on NeuroVault (https://neurovault.org/collections/11646/). The here used version of the Cyberball task can be shared upon request.

### 2.2 Participants

In all studies, NT participants were recruited by means of poster and online advertisements as well as an online participant recruitment system of the University of Vienna. Participants indicated via a thorough online questionnaire that they had no past or present psychiatric or neurological illnesses (NT sample only), had not taken part in similar studies before, had never studied psychology and had no risk factors concerning MR scanning (the latter criterion for the fMRI study only). In the fMRI study, ASD participants were recruited via an existing database of participants that registered in the past for taking part in university studies, as well as by contacting institutions that diagnose and treat people with ASD in Vienna and Linz. For all ASD participants, we confirmed a clinical ICD-10/DSM-IV diagnosis of the Asperger syndrome (F84.5/299.80) given by an accredited institution, and if possible assessed with the Autism Diagnostic Observation Schedule (Lord et al., 2000). Twenty-one ASD participants were matched to 21 NT participants regarding gender, age, handedness, and intelligence (see Table 1 for sample characteristics and matching criteria). Six ASD participants indicated having comorbidities such as depression or anxiety. A post hoc sensitivity power analysis in G*Power (version 3.1.9.2; Faul et al., 2007) showed that with our study we could have reasonably been able to detect a minimum effect size of Cohen’s *d* = 0.40 (α = 0.05, 1-β = 0.80).

**Table 1.** Sample characteristics and matching criteria in the fMRI study.

|  | NT | ASD |
| --- | --- | --- |
| **Nationality** |  |  |
| Austria | 16 | 19 |
| Germany | 5 | 2 |
| **Gender\*** |  |  |
| male | 16 | 15 |
| female | 5 | 6 |
| **Medication intake\*\*** |  |  |
| yes | 1 | 7 |
| no | 20 | 14 |
| **Handedness\*** |  |  |
| right | 20 | 20 |
| left | 1 | 1 |
| **Age\*** | 36.14 (10.16) | 36.52 (10.07) |
| **Intelligence\*** |  |  |
| SPM | 7.24 (1.79) | 7.62 (1.28) |
| MWT-B | 30.00 (4.34) | 29.57 (4.00) |

*Note.* Sociodemographic data of the participants in the fMRI study. Frequencies regarding nationality, gender, and handedness as well as averages (standard deviations) for age and intelligence are given. \*Matching criteria; \*\*The NT participant took thyroid and blood pressure medication, six autism spectrum disorder (ASD) participants took medication against psychiatric disorders such as e.g., depression, anxiety, or panic disorder, and one ASD participant took hormonal birth control; SPM = Standard Progressive Matrices; MWT-B = Mehrfachwahl-Wortschatz-Intelligenztest (version B).

### 2.3 Questionnaires

For matching purposes, two short forms of measures targeting general intelligence were given to all participants in the fMRI study. The Standard Progressive Matrices (SPM; Kratzmeier & Horn, 1988) is a nonverbal measure to assess general intelligence with figural material. The version used here contained nine items in each of which participants had to select the missing piece to an incomplete pattern. The Mehrfachwahl-Wortschatz-Intelligenztest (MWT-B; Lehrl, 1995) measures crystallized intelligence. For each item, participants needed to choose which one out of five given words is an existing word. We further assessed the following personality questionnaires: The degree to which a person reports traits associated with the autistic spectrum were measured using the 33-item version of the Autism-Spectrum-Quotient (AQ-k; Freitag et al., 2015). Those traits are based on the triad of impairments (impaired social communication, impaired social interaction and restricted, repetitive behaviors or activities) and on other areas of cognitive abnormality. They are viewed as part of a quantitative continuum where a person can be located based on their score. The cut-off score for a clinically significant level of autistic traits is ≥ 17. Alexithymia, the subclinical inability to identify and describe emotions, was measured with the 20-item version of the Toronto-Alexithymia-Scale (TAS-20; Bach et al., 1996; Bagby et al., 1994) and is divided into three subscales (Difficulty Identifying Feelings = DIF, Difficulty Describing Feelings = DFF, Externally-Oriented Thinking = EOT). To measure the subjective empathic qualities in the participants, the Interpersonal-Reactivity-Index (IRI) was used (Davis, 1980; Paulus, 2012). This multidimensional self-report questionnaire incorporates affective as well as cognitive aspects of the empathic reaction and is divided into four subscales, three measuring aspects of affective (Fantasy, Empathic Concern and Personal Distress) and one measuring cognitive empathy (Perspective Taking). Finally, the revised version of the Beck-Depression-Inventory (BDI-II; Kühner et al., 2007) was used to measure the amount of depressive symptoms. All questionnaires were administered via Paper & Pencil format in their German versions. Cohen’s *d*’s for questionnaire comparisons were calculated using the effect size calculation spreadsheet (version 4.2) provided by Lakens (2013).

### 2.4 Procedure

In all studies, each participant gave written consent at the outset of their respective session. The overall project was approved by the ethics committee of the Medical University of Vienna (EK 1166/2015) and performed in line with the latest revision of the Declaration of Helsinki (2013) on ethical principles for medical research involving human participants. For the fMRI study, each participant was asked not to take any medication, drugs, or alcohol 24 hours prior to the appointment. The task procedure was changed slightly to create a social situation adapted to the scanner environment. Participants came together with four confederates (always two males and two females) who were presented as other participants, but in fact were helpers of the experimenters merely acting as participants. This was done to make the social situation of the virtual ballgame more realistic and to make the participants believe that they were playing the game together with other participants. The five participants were given verbal instructions and played two short practice rounds on computers in the control room of the MRI scanner. The confederates were asked to remain seated and told they would be playing the main game from outside the scanner, while the participant was led into the scanner room. After general adjustments, each participant played three runs of the Cyberball game. After the task, the structural scan was acquired. When participants left the scanner room, they were asked to fill out additional questionnaires and told that the other participants had already left. Before leaving, each participant received 25 Euros as compensation and was fully debriefed by the experimenters about any deception used in the study, specifically to avoid negative emotions regarding exclusion by other individuals in the game. The whole procedure lasted around 2.5 hours.

### 2.5 Cyberball task

Participants played a modified version of Cyberball, a virtual computer-based ball-tossing game (Abrams et al., 2011; Eisenberger et al., 2003; Williams et al., 2000; Williams & Jarvis, 2006; Zadro et al., 2004). We created an adapted version fundamentally similar to the version used in Novembre et al. (2015), but using videos depicting white silhouettes of real humans on a black background^1^. Importantly, this version was designed to be used in the autistic population, ensuring as little distraction as possible by avoiding for example facial expressions and uneven background (see Figure 1). During the game, four avatars supposedly controlled by participants interacted in a virtual environment by throwing a ball to each other. The three main conditions differed in respect to whether: a) the participant was actively involved in the game (*self active, other active*) or only observing the others play (*other passive*), and b) whether the participants were asked to rate their own feelings (*self active*) or those of the player standing opposite of them (*other active, other passive*). Every condition included 16 trials of ball-tossing with approximately 13-15 ball throws per trial. During inclusion trials, the participant (the self) receives the ball and can throw it back to one of the others. Throwing was realized by pressing the arrow keys in the behavioral studies or the buttons on an fMRI-compatible button-box in the fMRI study. During exclusion trials, the player starts in the first trial with throwing the ball but then does not receive it back for the rest of that trial. Furthermore, the congruency to the emotional state of the (spatially) opposite person was varied via inclusion or exclusion of self and other in the game^2^.

**Figure 1.**
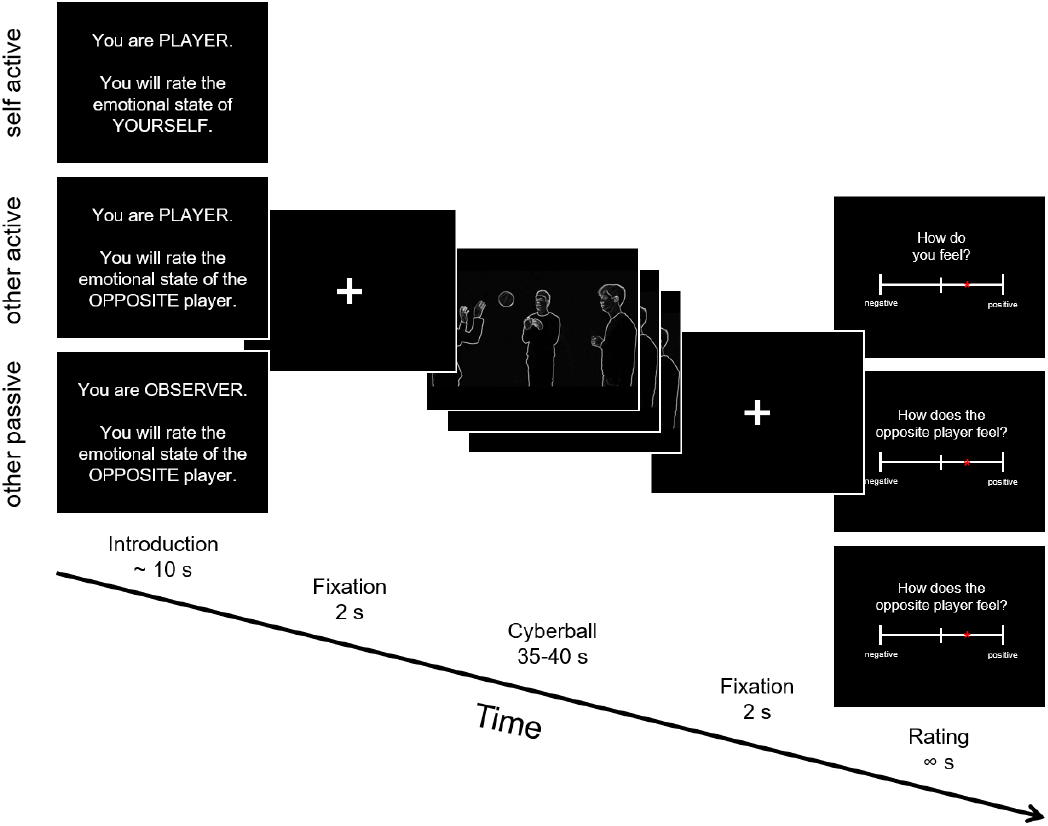
Schematic depiction of the Cyberball trial structure. Participants were shown an introduction screen with information whether they would be an active player or observer in the next game and whether they would judge the emotional state of themselves or of the player standing opposite of them. Following that was the Cyberball gameplay between 35 and 40 seconds, where different movie clips were shown depending on the left and right ball-throws of the participants. After each trial, emotional state ratings were collected.

During incongruent trials, one was excluded from the game while the other was excluded (i.e., self and other experiencing different emotions, either Self Included/Other Excluded or Self Excluded/Other Included). To measure self- and other-related emotional responses after each trial, participants were asked to rate either their own feelings (*self active*) or those of the person standing opposite of them (*other active* and *other passive*) on a continuous rating scale from negative to positive (representing exact values between -250 and +250, later rescaled to values between -10 and +10), staying on the screen until the response was acquired. The game was played in a fixed, previously pseudorandomized trial order in all studies and all participants. In the fMRI study, the three conditions were played in the scanner in separate runs, in one of six pseudorandomized orders. All videos shown in the game were filmed with a Panasonic Lumix DMC-FZ200 and later edited with Windows Movie Maker (Microsoft; applied setting: edge detection) and Corel VideoStudio Pro X8 (https://www.videostudiopro.com/en/pages/videostudio-x8/; applied filters: invert and monochrome) to a final resolution of 640×360 pixel. The Cyberball task was implemented in MATLAB 2010b (Mathworks, 2010), using Cogent 2000 (Version 1.29, http://www.vislab.ucl.ac.uk/cogent_2000.php).

### 2.6 fMRI Data acquisition

Functional data of the fMRI study was acquired using a 3 Tesla Siemens Magnetom Skyra MRI-system (Siemens Medical, Erlangen, Germany) in Vienna, equipped with a 32-channel head coil as well as a high-performance gradient system for fast, high-resolution whole brain multiband echoplanar imaging. The scanning sequence included the following parameters: Echo time (TE)/repetition time (TR) = 34/704 ms, flip angle = 50°, interleaved multi-slice mode, interleaved acquisition, field of view = 210 mm, matrix size = 96 × 96, voxel size = 2.2 × 2.2 × 3.5 mm^3^, 32 axial slices coplanar the connecting line between anterior and posterior commissure, and slice thickness = 3.5 mm. For each of the three conditions, functional volumes were acquired within 11-13 minutes durations per run (the exact number of volumes depending on the choice of ball throws and the rating velocity of the participants), with small breaks in between the runs. Structural images were acquired using a magnetization-prepared rapid gradient-echo sequence (TE/TR = 2.29/2300 ms, ascending acquisition, single shot multi-slice mode, 176 sagittal slices, voxel size = 0.9 × 0.9 × 0.9 mm^3^, flip angle = 8°, slice thickness = 0.94 mm, field of view = 240 mm).

### 2.7 Statistical analysis

The manuscript was checked for reporting errors in the statistical analyses using statcheck (http://statcheck.io/, Rife et al., 2016) and for reference errors using reciteworks (4cite Labs, https://reciteworks.com/).

#### Behavioral data

For the behavioral analyses, our aim was to investigate EAB and EEB as well as possible group differences in the fMRI study. Subjective ratings pertaining to the different conditions were entered in an analysis of variance (ANOVA) using the R package ezANOVA including the within-subjects factors *target* (self active, other active, other passive), *congruence* (congruent, incongruent), and *valence* (positive, negative), and the between-subjects factor *group* (ASD, NT). As in Silani et al. (2013), ratings in the unpleasant conditions were multiplied by -1 to allow the comparison of the bias across valences. All behavioral analyses were done in RStudio (version 4.1.0; R Core Team, 2020).

#### Brain data

##### Preprocessing and analysis

To preprocess and statistically analyze the fMRI data, the current version of the software Statistical Parametric Mapping (SPM12, Wellcome Trust Centre for Neuroimaging, https://www.fil.ion.ucl.ac.uk/spm/software/spm12/) running on MATLAB Version R2010b (Mathworks, 2010) was used. Preprocessing included realignment, coregistration of structural and functional images, segmentation into gray matter, white matter and cerebrospinal fluid, spatial normalization, and spatial smoothing by convolution with an 8 mm full-width at half-maximum (FWHM) Gaussian kernel. The first-level design matrix of each subject contained two separate regressors (playing time during trial and rating) for each of the four conditions (Self Included/Other Included, Self Included/Other Excluded, Self Excluded/Other Excluded, Self Excluded/Other Included), leading to a total of eight regressors for each of the three targets (self active, other active, other passive) and 24 regressors in total. In general, the full playing time of the ball game in each trial was modeled in a block-design fashion and convolved with SPM12’s standard canonical hemodynamic response function. Nuisance regressors included six realignment parameters and an additional two modeling (mean-centered) white matter and cerebrospinal fluid signal for each run (the latter two were extracted using the REX toolbox by Duff et al., 2007). For the group level, we created three contrast images incorporating the three conditions by contrasting congruent and incongruent trials and averaging over valence: EAB_active_ = self active (incongruent > congruent), EEB_active_ = other active (incongruent > congruent), and EEB_passive_ = other passive (incongruent > congruent). This averaging over valence for all analyses of the fMRI data was done purposefully as the stimuli shown for each valence (inclusion and exclusion) differed in important aspects such as motor movement and activity which would have influenced brain activity. For example, participants were only able to conduct one ball throw during exclusion trials while they were constantly throwing the ball in inclusion trials. This made an interpretation of the contrasts involving valence difficult.

##### Whole brain analysis

For the whole-brain analysis, the second-level model included a flexible-factorial design with the within-subject factors *subject* and *condition*, and the between-subjects factor *group* (NT vs. ASD). As a proof of concept and to evaluate whether our manipulation of congruency was successful, we first report the mean contrasts for each of the three conditions over both groups. To test our main hypotheses of increased EAB and EEB in the participants with ASD, we then report the group comparisons for each of the three conditions (EAB_active/NT_ vs. EAB_active/ASD_, EEB_active/NT_ vs. EEB_active/ASD_, EEB_passive/NT_ vs. EEB_passive/ASD_, as well as the complementary contrasts for ASD vs. NT). Finally, we report group comparisons for EEB_active_ after subtracting either EAB_active_ (as done in the previous study by Silani et al., 2013) or EEB_passive_ (our new control condition) as well as the additional contrast in our region-of-interest (ROI) analyses (see below): EEB_active_ + EEB_passive_ vs. EAB_active_, both for NT vs. ASD and vice versa. All whole brain imaging results are reported at a cluster probability of *p* < .05 (familywise-error (FWE)-corrected, cluster-forming threshold of *k* = 279, initial cluster-defining threshold *p* < .001 uncorrected). Anatomical regions were labelled with SPM12’s Anatomy toolbox (version 2.2c; Eickhoff et al., 2005).

##### Region of Interest analyses

Lastly, to explore differences in two major regions identified as being involved in social cognition, self-other distinction, and emotional egocentricity, we conducted ROI analyses in rSMG and rTPJ. We created 8mm spheres around two peak MNI coordinates taken from the independent study that also investigated self-other distinction in participants with and without ASD using the visuo-tactile egocentricity paradigm (Hoffmann, Koehne, et al., 2016): rSMG: x = 65, y = -37, z = 33; rTPJ: x = 51, y = -52, z = 21, using MarsBaR for ROI creation (Brett et al., 2002). We then extracted parameter estimates for each of the two ROIs using the first-level contrast images of each participant, separate for congruence and target (averaging over valence) with REX (Duff et al., 2007). We then calculated an ANOVA including the within-subjects factors *target* (self active, other active, other passive), *congruence* (congruent, incongruent), *roi* (rSMG, rTPJ) and the between-subjects factor *group* (ASD, NT). We focused these analyses on two aspects inherent in our task: 1) Checking for group differences in emotional egocentricity related to the degree of emotional involvement by contrasting the other active and other passive condition, and 2) checking for differences in ego-vs. altercentric judgments by contrasting all other-related (other active and other passive) with the self-related condition (self active). Mauchly’s test for sphericity was not statistically significant for any effects including the factor target, thus fulfilling the sphericity assumption.

## 3 Results

### Questionnaire results

In the personality trait questionnaires, the two groups differed significantly in their reported autistic, alexithymic and depressive traits, with ASD subjects consistently indicating higher values than NT (see Table 2 for an overview). Regarding empathic abilities, ASD participants described significantly greater personal distress, while NT ascribed themselves significantly higher perspective taking abilities. The two groups did not significantly differ regarding their trait empathic concern and fantasy.

**Table 2.** Differences between ASD and NT participants in trait personality questionnaires in the fMRI study.

|  | NT | ASD | *t* | *df* | *p* | Cohen's *d* |
| --- | --- | --- | --- | --- | --- | --- |
| **Autistic traits** | 6.76 (4.48) | 23.95 (5.62) | -10.96 | 40 | < .001 | 3.47 |
| **Alexithymia** |  |  |  |  |  |  |
| DIF | 12.29 (4.45) | 20.14 (5.70) | -4.98 | 40 | < .001 | 1.57 |
| DDF | 11.47 (3.68) | 17.43 (4.04) | -4.99 | 40 | < .001 | 1.58 |
| EOT | 16.09 (4.11) | 20.52 (4.86) | -3.19 | 40 | .003 | 1.01 |
| **Empathy** |  |  |  |  |  |  |
| Fantasy | 3.11 (0.68) | 2.68 (0.85) | 1.80 | 40 | .079 | 0.57 |
| Empathic Concern | 3.66 (0.66) | 3.37 (0.76) | 1.33 | 40 | .189 | 0.42 |
| Personal Distress | 2.51 (0.43) | 3.09 (0.79) | -2.99 | 40 | .004 | 0.93 |
| Perspective Taking | 3.69 (0.49) | 3.04 (0.66) | 3.61 | 40 | < .001 | 1.15 |
| **Depression** | 6.24 (6.30) | 13.29 (10.01) | -2.73 | 40 | .009 | 0.86 |

*Note.* Trait questionnaire data of the fMRI study separated by group. Average item sum scores (autism, alexithymia, depression) or average item scores (empathy) plus standard deviations in brackets. DIF = Difficulty Identifying Feelings subscale, DDF = Difficulty Describing Feelings subscale, EOT = Externally Oriented Thinking subscale.

### Behavioral task results

The analysis of the pilot study data can be found in the Supplementary Material (Tables S1 and S2). These analyses revealed conclusive evidence that our version of the Cyberball task was able to manipulate the congruency of simultaneous emotional states, as we observed an EEB and EAB in the participants’ ratings in pilot study 1, indexed by significant main effects of congruence and target x congruence interactions (all *p*’s ≤ .001). Furthermore, we established the third condition, other passive, as a valid control condition for the EEB, as no difference between congruent and incongruent condition (i.e., no bias) was observed there in pilot study 2 (see also Figure 2A and 2B)^3^.

**Figure 2.**
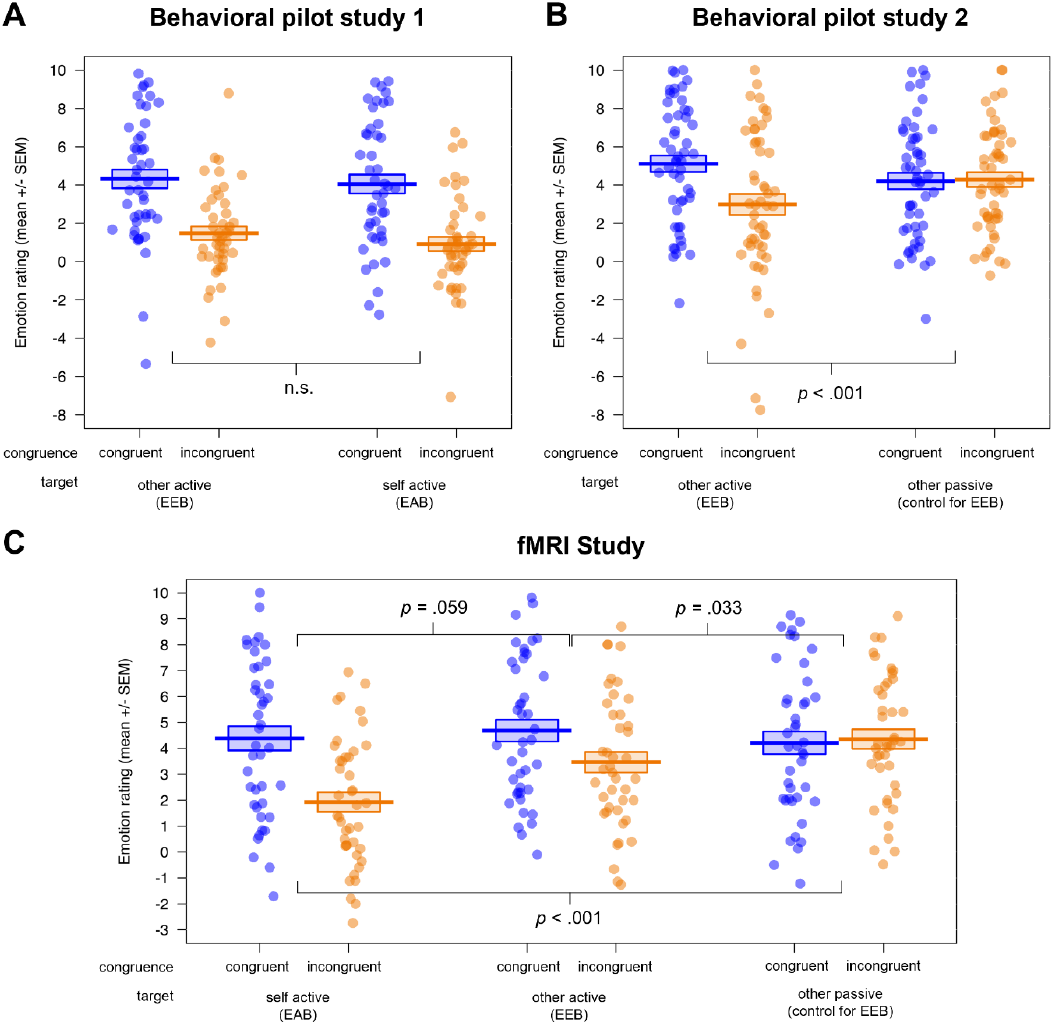
Visualization of the behavioral results in A) pilot study 1 (*n* = 45), B) pilot study 2 (*n* = 52) and C) the fMRI study (*n* = 42). All studies showed a successful induction of emotional biases (EEB and/or EAB), visible in the rating differences for congruent vs. incongruent emotional states. Additionally, pilot study 2 and the fMRI study validated the passive viewing condition as a control condition for the EEB. EEB = emotional egocentricity bias; EAB = emotional altercentricity bias.

In the ANOVAs of the fMRI study rating data (see Table S3 in the Supplementary Material), we observed main effects of congruence (*F*(1,40) = 18.06, *p* < .001, *η*^*2*^ = 0.031) and valence (*F*(1,40) = 4.57, *p* = .039, *η*^*2*^ = 0.021), with more extreme ratings for congruent (*M* ± *SD* = 4.43 ± 3.49) compared to incongruent (*M* ± *SD* = 3.25 ± 3.47) emotional states, and for negative (*M* ± *SD* = 4.31 ± 3.55) compared to positive (*M* ± *SD* = 3.36 ± 3.44) emotional states of the person to be judged. Furthermore, a main effect of target (and follow-up post hoc comparisons with the function pairs) indexed that the ratings in the self active (*M* ± *SD* = 3.16 ± 3.68) differed significantly from those in the other active (*M* ± *SD* = 4.07 ± 3.53; *p* = .044) as well as the other passive condition (*M* ± *SD* = 4.29 ± 3.26; *p* = .009), independent of congruence or valence (*F*(2,80) = 13.36, *p* < .001, *η*^*2*^ = 0.022).

A significant target x congruence (*F*(2,80) = 13.43, *p* < .001, *η*^*2*^ = 0.026; and follow-up post hoc comparisons with the function pairs) showed that the difference in congruent vs. incongruent ratings, i.e. the emotional bias, was significantly lower for the other passive (control for EEB; *M*_*diff*_ = 0.14) compared to the self active (EAB; *M*_*diff*_ = -2.46; *p* < .001) as well as compared to the other active condition (EEB; *M*_*diff*_ = -1.22; *p* = .033). Mirroring the results of pilot study 1, there was no significant difference between EAB and EEB (*p* = .059). We also observed a group x target interaction (*F*(1,40) = 3.24, *p* = .044, *η*^*2*^ = 0.005), showing that individuals with ASD had more extreme ratings compared to NTs in the self active (*M*_*diff*_ = 1.68; *p* = .021), but not in the other active (*M*_*diff*_ = 0.64; *p* = .535) or the other passive condition (*M*_*diff*_ = -0.06; *p* = .925). Lastly, we found a target x congruence x valence interaction (*F*(2,80) = 4.85, *p* = .010, *η*^*2*^ = 0.004), which showed that the rating difference between congruent and incongruent as well as positive and negative emotional states was highest for self active (*M* ± *SD* = 2.08 ± 4.08), medium for other passive (*M* ± *SD* = 0.99 ± 4.52) and lowest for other active (*M* ± *SD* = -0.03 ± 5.18). Importantly, an absence of any other group effects and specifically no significant congruence x group or congruence x target x group interactions (all *p*’s > .075) showed that both groups were equally susceptible to emotional biases on the behavioral level.

### Whole brain results

In our whole brain analyses, we evaluated the three target contrasts over both groups (see Figure 3 here and Tables S4-S6 in the Supplementary Material). In the mean contrast for self active (EAB) over both groups, we observed, among others, increased hemodynamic activity in bilateral middle temporal, frontal and superior medial gyri, right superior temporal gyrus, right superior parietal lobule, bilateral angular gyrus, right precuneus and right inferior frontal gyrus. Activity in these regions was increased during incongruent compared to congruent trials, when participants played the game and were asked to judge their own emotional state.

**Figure 3.**
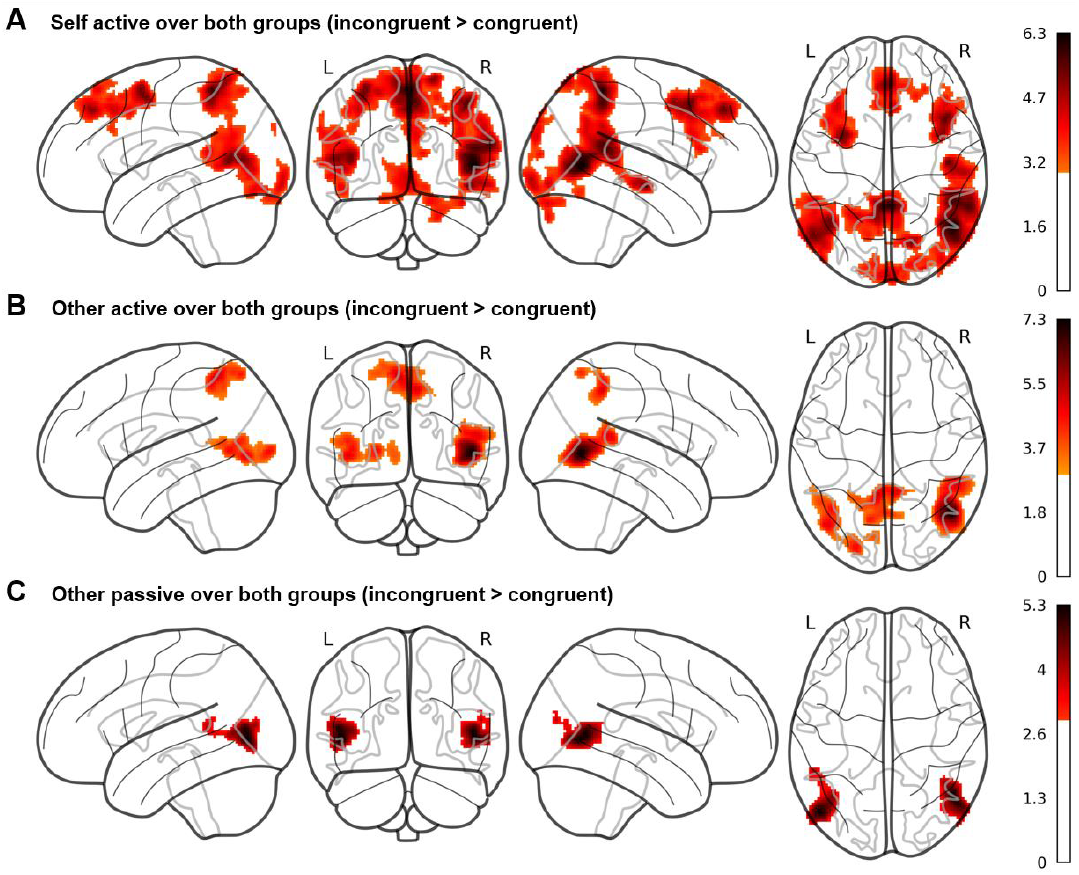
Whole brain results of the fMRI study for the three Cyberball conditions A) self active, B) other active, and C) other passive. Contrasts are averaged over the factors valence and group, calculated as incongruent > congruent conditions and displayed at a cluster probability of *p* < .05 (familywise-error (FWE)-corrected, cluster-forming threshold of *k* = 279, initial cluster-defining threshold *p* < .001 uncorrected). The results show activity in brain regions such as precuneus and superior temporal gyrus for the self active and other active conditions, while these regions are not active in the other passive condition.

In the mean contrast for other active (EEB) over both groups (incongruent compared to congruent trials), we observed increased hemodynamic activity within the right middle and superior temporal gyri, left middle occipital gyrus and bilateral precuneus including the left superior parietal lobule. Activity in these regions was increased during incongruent compared to congruent trials, when participants played the game but judged the other’s emotional state.

In the mean contrast for other passive over both groups, we observed increased hemodynamic activity in bilateral middle and left superior temporal gyri as well as middle occipital gyrus. Activity in these regions was increased during incongruent compared to congruent trials, when participants were again asked to judge the other’s emotional state but were passively watching the game as an observer, while four others were playing.

However, evaluating our main hypothesis for group differences in the self active, other active and other passive conditions (either the individual conditions or as a differential contrast for EEB_active_ – EAB_active_ or EEB_active_ – EEB_passive)_, we did not observe any increased hemodynamic activity in either of the two groups. In fact, none of our whole brain group comparisons showed any active clusters at an FWE-corrected threshold. As we were specifically interested in the differential roles of two specific brain regions previously shown to be involved in self-other distinction, rTPJ and rSMG, we went on to analyse extracted brain activity in these areas.

### Region of interest results

In our ROI analysis, evaluating group differences in extracted brain activity in rSMG and rTPJ, we observed a main effect of congruence, showing increased activity in incongruent (*M* ± *SD* = -0.02 ± 0.008) compared to congruent (*M* ± *SD* = -0.03 ± 0.007) situations, independent of the target, roi or group membership: *F*(1,40) = 16.89, *p* < .001, *η*^*2*^ = 0.005 (see Table S7 for the full ANOVA results). We also found a main effect of roi, showing increased activity for rTPJ (*M* ± *SD* = 0.02 ± 0.008) compared to rSMG (*M* ± *SD* = -0.06 ± 0.007), independent of target, congruence, or group membership: *F*(1,40) = 29.07, *p* < .001, *η*^*2*^ = 0.11. Next, we observed a congruence x roi interaction, showing that the difference in activity between incongruent compared to congruent situations was increased in rTPJ (incongruent: *M* ± *SD* = 0.03 ± 0.01, congruent: *M* ± *SD* = 0.004 ± 0.01) compared to rSMG (incongruent: *M* ± *SD* = -0.07 ± 0.009, congruent: *M* ± *SD* = 0.06 ± 0.009), independent of target or group membership: *F*(1,40) = 23.75, *p* < .001, *η*^*2*^ = 0.002. Interestingly, we found a significant four-way interaction (group x congruence x target x roi, *F*(2,80) = 5.45, *p* = .006, *η*^*2*^ = 0.001). To follow up on this interaction, we calculated two orthogonal Helmert contrasts, contrasting (1) the degree of emotional involvement during emotional egocentricity (other active - other passive), and 2) ego-vs altercentric conditions (0.5 * other active + 0.5 * other passive - self active) and compared the difference scores for incongruent – congruent situations for each roi between the two groups to follow up on the four-way interaction. This revealed that the difference in activity between other active and other passive conditions between NT and ASD group was increased in rTPJ compared to rSMG, with the NT group showing higher activity (*p* = .034; see Figure 4A). In contrast, the difference in activity between the other- and self-related conditions between NT and ASD group was increased in rSMG compared to rTPJ, with the NT group showing higher activity (*p* = .017; see Figure 4B). In other words, while group differences in rTPJ activity seem to underpin evaluating egocentric judgements during active emotional involvement, group differences in rSMG relate to other-compared to self-related judgements. In both cases, participants with ASD had decreased activity in these regions compared to NTs.

**Figure 4.**
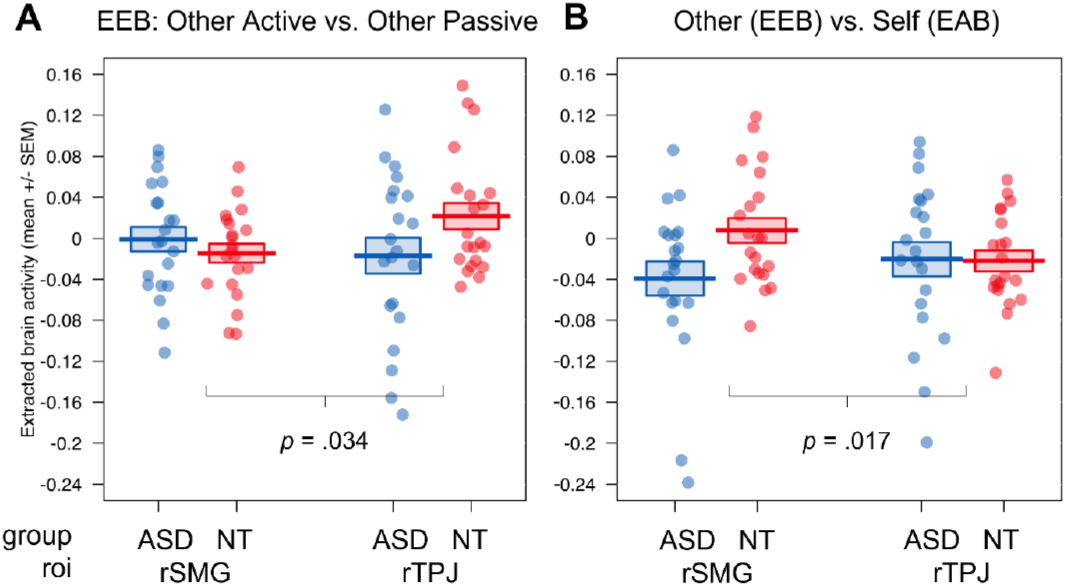
ROI results of the fMRI study for the group comparison of ASD vs. NT. Visualization of the four-way interaction group x congruence x target x roi, A) the comparison of emotional involvement during egocentric judgements (other active vs. other passive), B) the comparison of ego-(other active + other passive) vs. altercentric judgement conditions (self active); Displayed brain activity is averaged over the factors valence and calculated as incongruent – congruent; EEB = emotional egocentricity bias; EAB = emotional altercentricity bias; SEM = standard error of the mean, ASD = autism spectrum disorder, rTPJ = right temporoparietal junction, rSMG = right supramarginal gyrus.

## 4 Discussion

The present study investigated self-other distinction and the occurrence of ego- and altercentric biases in individuals with and without ASD.

Extending previous work on the topic (Hoffmann, Koehne, et al., 2016), we were particularly interested at evaluating egocentric and altercentric biases independently, rather than one against the other, as both biases reflect deficits in self-other distinction. To do this, we introduced an emotionally neutral condition, in which participants were asked to judge another person’s emotions without being involved in the situation themselves. This condition served to control for task complexity (i.e., incongruency between different emotional states) without cancelling out the processing of self-vs. other-related representations. Furthermore, we were interested to directly link behavioral group differences to differences in neural activation, by means of fMRI investigation. On the behavioral level, the two behavioral pilot studies as well as the fMRI study confirmed the validity of our newly adapted Cyberball version. In all three studies, when individuals were actively playing the game and had to deal with incongruent compared to congruent emotional states of self and other, (a) their judgment of the other person’s emotions was shifted towards their own emotional state (EEB), and (b) the judgment of their own emotions was shifted towards the other’s emotional state (EAB). The strength of these two biases was similar for pilot study 1 and the fMRI study.

Additionally, both pilot study 2 and the fMRI study showed that the EEB was only observable during active involvement in the game and not in the condition of passively observing the same game between four other players. In contrast to the two active conditions, passive observation led to more accurate, i.e., similar, judgments in both congruent and incongruent situations, possibly due to a lack of interference with one’s own emotional state. In sum, our new version of Cyberball was able to produce feelings of social inclusion/exclusion and generated emotional biases to a similar extent compared with other paradigms who reported affective biases in other younger and adult neurotypical samples inducing social inclusion/exclusion (Seidel et al., 2013) as well as for other emotions like envy or schadenfreude (Steinbeis et al., 2014; Steinbeis & Singer, 2014), pleasant or unpleasant visuo-gustatory (Hoffmann et al., 2015), visuo-tactile (Riva et al., 2016; Silani et al., 2013; Tomova et al., 2014), audio-visual (von Mohr et al., 2020) or face stimuli (Naor et al., 2018; Trilla, Eiserbeck, et al., 2020; Trilla, Weigand, et al., 2020). Our results therefore extend previous findings on emotional alter- and egocentric biases to the domain of ostracism. Looking at behavioural group differences, we could not confirm our hypothesis of stronger emotional altercentric or egocentric biases in the autistic compared to the NT population. Both biases were similarly high in the two groups, and significantly different from the passive condition, which is in line with the previous study indicating lack of self-other distinction impairments in autism (Hoffmann, Koehne, et al., 2016).

On the neural level, the ROI analyses in rTPJ and rSMG revealed differences depending on the region, target, and group. In particular, we observed increased brain activity in rTPJ but not rSMG in NTs compared to ASDs when actively dealing with the EEB (compared to passively watching the ballgame). This indicates that individuals with ASD recruit rTPJ to a lesser extent than NT individuals when suppressing their own emotions to accurately judge the emotional state of another person, while this is not the case for rSMG. We further observed increased brain activity in rSMG but not rTPJ in NTs compared to ASDs when dealing with ego-compared to altercentric judgements. A specific difference emerged when overcoming one’s own (EEB) compared to overcoming others’ (EAB) emotional states influence. Our results nicely complement and extend the findings reported in Hoffmann, Koehne, et al. (2016), who observed reduced connectivity of regions of the Theory of Mind network (rTPJ) but an unimpaired rSMG network connectivity during rest in individuals with ASD. In line with these findings, we show differential involvement of rTPJ and rSMG when dealing with conflicting emotional states in the same participant sample. This crucially underlines the necessity for a distinction between rSMG and rTPJ, and between ego-vs. altercentric biases, when investigating self-other distinction abilities in ASD. Future studies should focus on carefully teasing apart the separate roles of these two regions, for example, by employing more causal methods such as repetitive transcranial magnetic stimulation as in Silani et al. (2013).

Generally, the brain results also showed that our intended manipulation of congruency regarding self- and other-related emotional states was successful. When actively playing the game and either judging their own or the other’s incongruent emotional state, participants activated regions previously related to self-other distinction (Lamm et al., 2016; Steinbeis, 2016), self/other attribution (Kestemont et al., 2015), conflict monitoring (Jeannerod, 2018), mental imagery to represent the perspective of another person (Cavanna & Trimble, 2006) and theory of mind (especially false-belief reasoning) as well as visual perspective taking (see Molenberghs et al., 2016; Schurz et al., 2013, 2020 for meta-analyses). Mentalizing abilities seem to be required more strongly in situations where two emotional states are incongruent compared to congruent (Silani et al., 2013). This was not the case in the passive viewing condition, where participants merely observed other individuals playing the Cyberball game. The active conditions thus recruited more cognitive self-other distinction processes than the passive condition, where participants had no own emotional state to keep track of.

In contrast to previous research, we did not specifically find increased brain activity in rSMG during incongruent compared to congruent judgements in our whole brain analyses. However, it should be noted that the crucial role of the rSMG in overcoming the EEB was found in a visuo-tactile paradigm (Silani et al., 2013) and a task using monetary rewards and punishment (Steinbeis et al., 2014). Social emotions like in- or exclusion, on the other hand, belong to a different domain that could be processed in a different, possibly more cognitive way. Especially our task could have required more mentalizing abilities, as participants had to infer the emotional state of the other person who was displayed as an avatar, which might have recruited rTPJ more than rSMG.

We would also like to briefly discuss and address the major strengths and limitations of the present study. Firstly, it has to be acknowledged that the two groups were closely matched regarding handedness, gender, age, general intelligence, and differed significantly in their autistic traits, but also differed in their level of reported depressive traits. However, the higher levels of depression in the ASD group were very close to the cutoff for no to minimal depressive symptoms (Kühner et al., 2007). Furthermore, the ASD group reported increased alexithymic traits compared to the NT group. This is not surprising, as higher alexithymic symptoms in ASD have been reported in multiple studies and alexithymia often co-occurs in individuals with ASD (Bird et al., 2010; Bird & Cook, 2013; Bird & Viding, 2014; Chester et al., 2013). Including alexithymia as a covariate in our analyses did not change any of our here reported results. Nevertheless, as Bird et al. (2010) reported that the strength of left anterior insula activation in response to others’ suffering was predictive of the degree of alexithymia in both autistic and NT groups but did not in fact vary as a function of group, future studies should better tease apart the proneness to emotional ego- and altercentric biases by specifically recruiting both NT and autistic individuals with high and low levels of alexithymia.

Secondly, the study used an adapted version of Cyberball with videos depicting real-life figures instead of cartoon animations. This was done to increase ecological validity and develop a task suitable for use in the autistic population that did not contain distractions. The game avoided solving the task by using emotional facial expressions, as all avatars kept a neutral expression in the video and the videos were edited to show white silhouettes on black background. Furthermore, four confederates were introduced to the participant at the beginning of the session to create a realistic social situation and immerse the participants in the live gameplay. However, due to the length of the game and somewhat predictable nature of the conditions (i.e., the varying valence and congruence to measure EAB and EEB), it could well be that participants emotionally detached or disengaged from the game at some point and weren’t able to empathize as strongly. However, the results of an increased behavioral EAB and EEB in the active but not the passive conditions in all our three studies, combined with the recruitment of mentalizing regions in our fMRI study, showed that participants were still influenced by their own and the other person’s emotions when they were actively engaged in the game. And as described by Zadro et al. (2004), even ostracism by a computer compared to ostracism by real individuals is able to produce similar levels of social exclusion. Nevertheless, future studies using these types of social interaction setups should consider even stronger “interactive” playing modes that introduce greater variability and constantly remind the participant of the other players, e.g., via live videos.

Lastly, we did not include a passive condition for the altercentric bias, which could have made results regarding the EAB more informative. Future studies should therefore pay close attention to how the biases are being measured and include appropriate controls for all of them.

In conclusion, the present findings replicate previous behavioral and neurophysiological results on the ego- and altercentric biases in the emotional domain and expand them to the field of ostracism. Furthermore, they suggest no behavioral differences in the processing of simultaneous emotional states and thus intact self-other distinction in ASD, as well as specific neurophysiological differences rooted in rSMG and rTPJ between individuals with and without ASD in dealing with ego- and altercentric biases. This study has crucial implications for further research of social cognition abilities in ASD. Investigating socio- emotional competences on a more basic level using valid paradigms will ultimately pave the way to better understand individuals with ASD and could set a foundation for interventions promoting successful and long-lasting social interactions and relationships.

## Acknowledgements

We thank Roger Mundry for statistical guidance. We thank Seda Özoglu and Aaron Salzer for planning and conducting the two behavioral pilot studies together with PS and GS. Lastly, we would like to acknowledge the help of all confederates aiding in the fMRI study.

## CRediT author statement (Brand et al., 2015)

**HH:** Conceptualization, Methodology, Formal analysis, Investigation, Data Curation, Writing - Original Draft, Writing - Review & Editing, Visualization, Project administration, Funding acquisition; **LL:** Conceptualization, Formal analysis, Investigation, Writing - Original Draft, Writing - Review & Editing; **HHH:** Methodology, Investigation, Data Curation, Writing - Review & Editing, Funding acquisition; **PS:** Conceptualization, Methodology, Investigation, Data Curation, Writing - Review & Editing; **GS:** Conceptualization, Methodology, Software, Investigation, Resources, Writing - Original Draft, Writing - Review & Editing, Supervision, Project administration, Funding acquisition.

## Footnotes

1 The videos displayed male participants trained to show similar body posture during throwing and catching the ball, while the rhythm of the ball throws was clocked using a metronome. One person stood behind the camera catching and throwing the ball in front of the camera lens to create an egocentric perspective and viewed the other players standing to his/her left and right, and in front of the participant.

2 To achieve these conditions, participants were ostensibly divided into two groups and informed that while Group A would only be able to throw to the left and right player, Group B would be able to throw to the left, right and front players. Participants were purposely always assigned to Group A, together with the opposite player. The deception was used to avoid participants throwing the ball to the opposite player after seeing him excluded by the other two.

3 Note that in each pilot study only one control condition (either self active in pilot 1 or other passive in pilot 2) was implemented. In the fMRI study, both control conditions were employed.

