## Supplementary Material for "Emotional ego- and altercentric biases in high-functioning autism spectrum disorder: Behavioral and neurophysiological evidence"

[#] Shared first authorship

* Corresponding authors:, +43-1-4277-47223, Department of Clinical and Health Psychology, University of Vienna, Liebiggasse 5, 1010 Vienna, Austria;, +43-1-4277-47130, Social, Cognitive and Affective Neuroscience Unit, Department of Cognition, Emotion, and Methods in Psychology, University of Vienna, Liebiggasse 5, 1010 Vienna, Austria.

### S.1 Behavioral pilot studies

*Sample and Procedure*. The aim of behavioral pilot study 1 was to validate the newly created Cyberball paradigm using the same structure of the original visual-tactile paradigm (Silani et al., 2013). Behavioral pilot study 2 subsequently aimed at introducing an additional control condition for the EEB. In both pilots, the participants came in groups of 8-10 to a computer room at the university. Participants were instructed thoroughly before the start of the task and had the chance to ask and clear any questions. They were instructed to sit in a circle, meant to create a social situation, and told that they would play a virtual ball-tossing task with each other. The task procedure was very similar in both behavioral studies, with the only difference that in pilot 1, participants were asked to come in pairs of two and bring a friend of the same gender, who they would (allegedly) be playing with in the ballgame. This procedure was dropped in the second pilot and the fMRI study for easier recruitment purposes and believability of the cover story. In pilot 2, participants were told they would be connected randomly with the other participants via computers. Participants were then randomly assigned to one of the computers in the experimental room. In general, brief trainings of each game round were played before the actual game. It was ensured that all participants started the game at the same time to ensure believability of the virtual connection and interaction in the game. In reality, each participant played against the computer and completed pre-specified trials measuring the EAB and EEB in different conditions. After playing, participants were debriefed about all deceptive elements of the study. The whole procedure lasted approximately one hour, and each participant received a compensation of 10€. Pilot study 1 included 45 participants (26 females and 19 males, age range 18-55 years), pilot study 2 included 52 participants (24 females and 28 males, age range 20-41 years).

*Results*. As in the fMRI study, we calculated one analysis of variance (ANOVA) for each pilot study, including the within-subjects factors *target* (pilot 1: self active and other active; pilot 2: other active and other passive), *congruence* (congruent, incongruent) and *valence* (positive, negative). Here we report the ANOVA tables of the rating data in the two behavioral pilot studies 1 (Table S1) and 2 (Table S2). In pilot study 1, we observed main effects of congruence ($F$(1,44) = 55.88, $p$ < .001, $\eta^2$ = 0.131) and valence ($F$(1,44) = 6.02, $p$ = .018, $\eta^2$ = 0.039), showing that ratings were more extreme for congruent ($M \pm SD$ = 4.20 ± 4.51) compared to incongruent ($M \pm SD$ = 1.23 ± 4.49) emotional states, and for negative ($M \pm SD$ = 3.50 ± 4.48) compared to positive ($M \pm SD$ = 1.94 ± 4.87) emotional states of the person to be judged. Especially the main effect of congruence demonstrates that our sample indeed showed emotional biases during our version of the Cyberball task. Furthermore, a congruence x valence interaction ($F$(1,44) = 12.15, $p$ = .001, $\eta^2$ = 0.018) showed that the rating difference between incongruent and congruent emotional states, i.e., the strength of the bias, was higher for when the evaluated person was in a positive ($M_{diff}$ = 3.96) compared to a negative ($M_{diff}$ = 1.98) emotional state, independent of whether that person was oneself or the other. In sum, pilot study 1 demonstrated significant emotional biases and a significantly stronger EAB compared to EEB. Lastly, the non-significant target x congruence interaction showed that the rating difference between congruent and incongruent emotional states, i.e. the bias, was similar for self active ($M_{diff}$ = 3.12) compared to other active ($M_{diff}$ = 2.82), independent of the valence of the rated emotional state.

Table S1

*ANOVA using the rating data of the behavioral pilot study 1.*

| Effect | $\hat{\eta}^2_G$ | 90% CI | $F$ | $df$ | $df_{\mathrm{res}}$ | $p$ |
|---|---|---|---|---|---|---|
| Target | .002 | [.000, .068] | 2.15 | 1 | 44 | .150 |
| Congruence | .131 | [.016, .292] | 55.88 | 1 | 44 | < .001 |
| Valence | .039 | [.000, .168] | 6.02 | 1 | 44 | .018 |
| Target × Congruence | .001 | [.000, .042] | 0.57 | 1 | 44 | .453 |
| Target × Valence | .003 | [.000, .078] | 3.75 | 1 | 44 | .059 |
| Congruence × Valence | .018 | [.000, .128] | 12.15 | 1 | 44 | .001 |
| Target × Congruence × Valence | .000 | [.000, .000] | 0.01 | 1 | 44 | .921 |

As the EAB was bigger than the EAB and thus not an optimal control condition as used in our previous study (Silani et al., 2013), we conducted a second pilot study. There, we compared EEB and a new control condition of passive viewing and we again observed, in line with our expectations, a main effect of congruence ($F$(1,51) = 11.28, $p$ = .001, $\eta^2$ = 0.017), whereby ratings were more extreme for congruent ($M \pm SD$ = 4.69 ± 4.28) compared to incongruent ($M \pm SD$ = 3.65 ± 5.02) emotional states. This is evidence for an EEB in this sample. Interestingly, we also observed a target x congruence interaction ($F(1,51) = 19.43$, $p < .001$, $\eta^2 = 0.002$), showing that the rating difference between congruent and incongruent emotional states, i.e., the EEB, was higher for other active ($M_{diff}$ = 2.12) compared to other passive ($M_{diff}$ = -0.05), independent of the valence of the rated emotional state. In sum, pilot study 2 clearly showed that while an EEB was present in this sample during active playing of the game, it disappeared in the condition of passive viewing (where the ratings for congruent and incongruent emotional states of two other players were nearly identical).

Table S2

*ANOVA using the rating data of the behavioral pilot study 2.*

| Effect | $\hat{\eta}^2_G$ | 90% CI | $F$ | $df$ | $df_{\text{res}}$ | $p$ |
|---|---|---|---|---|---|---|
| Target | .001 | [.000, .041] | 0.49 | 1 | 51 | .487 |
| Congruence | .017 | [.000, .116] | 11.28 | 1 | 51 | .001 |
| Valence | .008 | [.000, .091] | 1.75 | 1 | 51 | .192 |
| Target × Congruence | .018 | [.000, .119] | 19.43 | 1 | 51 | < .001 |
| Target × Valence | .000 | [.000, .000] | 0.00 | 1 | 51 | .991 |
| Congruence × Valence | .002 | [.000, .062] | 1.88 | 1 | 51 | .177 |
| Target × Congruence × Valence | .000 | [.000, .000] | 0.02 | 1 | 51 | .879 |

### S.2 fMRI study

*Behavioral results*. Table S3 shows the ANOVA table of the rating data in the fMRI study.

Table S3

*ANOVA using the rating data of the fMRI study.*

| Effect | $\hat{\eta}^2_G$ | 90% CI | $F$ | $df$ | $df_{\text{res}}$ | $p$ |
|---|---|---|---|---|---|---|
| Group | .025 | [.000, .149] | 2.45 | 1 | 40 | .126 |
| Target | .022 | [.000, .085] | 13.36 | 2 | 80 | < .001 |
| Congruence | .031 | [.000, .163] | 18.06 | 1 | 40 | < .001 |
| Valence | .021 | [.000, .141] | 4.57 | 1 | 40 | .039 |
| Group × Target | .005 | [.000, .037] | 3.24 | 2 | 80 | .044 |
| Group × Congruence | .000 | [.000, .000] | 0.05 | 1 | 40 | .831 |
| Group × Valence | .002 | [.000, .072] | 0.46 | 1 | 40 | .503 |
| Target × Congruence | .026 | [.000, .092] | 13.43 | 2 | 80 | < .001 |
| Target × Valence | .001 | [.000, .004] | 0.80 | 2 | 80 | .453 |
| Congruence × Valence | .006 | [.000, .098] | 3.08 | 1 | 40 | .087 |
| Group × Target × Congruence | .001 | [.000, .002] | 0.70 | 2 | 80 | .500 |
| Group ×Target × Valence | .002 | [.000, .004] | 0.99 | 2 | 80 | .374 |
| Group × Congruence × Valence | .006 | [.000, .100] | 3.35 | 1 | 40 | .075 |
| Target × Congruence × Valence | .004 | [.000, .030] | 4.85 | 2 | 80 | .010 |

| | | | | | | |
|---|---|---|---|---|---|---|
| Group × Target × Congruence × Valence | .001 | [.000, .001] | 1.52 | 2 | 80 | .225 |

*Whole brain results*. Here we report the whole brain results of the fMRI analyses, namely for the contrasts self active (incongruent > congruent) in Table S4, other active (incongruent > congruent) in Table S2 S5 and other passive (incongruent > congruent) in Table S6, all averaged over both groups.

Table S1

*fMRI whole brain results for the contrast self active (incongruent > congruent).*

| Clusters and brain regions | h | *x* | *y* | *z* | *t* value | *p* value |
|---|---|---|---|---|---|---|
| **Cluster 1** (*k* = 5724) | | | | | | < .001 |
| Middle temporal gyrus | R | 50 | -64 | 4 | 6.33 | |
| Middle temporal gyrus | R | 50 | -46 | 16 | 5.70 | |
| Superior temporal gyrus (Inferior parietal lobule) | R | 60 | -48 | 18 | 5.61 | |
| **Cluster 2** (*k* = 3085) | | | | | | < .001 |
| Precuneus | R | 2 | -48 | 58 | 6.05 | |
| Precuneus | R | 4 | -48 | 50 | 5.69 | |
| Superior parietal lobule | R | 18 | -76 | 66 | 5.25 | |
| **Cluster 3** (*k* = 1486) | | | | | | < .001 |
| Middle frontal gyrus | R | 44 | 16 | 50 | 5.48 | |
| Precentral gyrus | R | 40 | 4 | 50 | 5.15 | |
| Inferior frontal gyrus (p. triangularis) | R | 38 | 18 | 26 | 4.19 | |
| **Cluster 4** (*k* = 1436) | | | | | | < .001 |
| Superior medial gyrus | R | 2 | 40 | 44 | 5.43 | |
| Posterior-medial frontal | R | 6 | 24 | 54 | 4.42 | |
| Superior medial gyrus | L | -4 | 36 | 60 | 4.24 | |
| **Cluster 5** (*k* = 2707) | | | | | | < .001 |
| Middle temporal gyrus | L | -52 | -70 | 8 | 5.39 | |
| Middle temporal gyrus | L | -42 | -60 | 8 | 5.39 | |
| Middle temporal gyrus | L | -52 | -54 | 4 | 4.27 | |
| **Cluster 6** (*k* = 1150) | | | | | | < .001 |
| Middle frontal gyrus | L | -32 | 2 | 56 | 5.36 | |
| Middle frontal gyrus | L | -36 | 24 | 44 | 4.47 | |
| Middle frontal gyrus | L | -42 | 12 | 44 | 4.08 | |

*Note.* Significant clusters including hemisphere h, cluster size *k*, MNI coordinates x, y, z, *t* value and *p* value (whole brain, FWE-corrected at $p < .05$, cluster level, $k > 279$).

Anatomical regions were labelled with SPM12's Anatomy toolbox (version 2.2c; Eickhoff et al., 2005)

Table S2

*fMRI whole brain results for the contrast other active (incongruent > congruent).*

| Clusters and brain regions | h | *x* | *y* | *z* | *t* value | *p* value |
|---|---|---|---|---|---|---|
| **Cluster 1** (*k* = 1650) | | | | | | < .001 |
| Middle temporal gyrus | R | 48 | -64 | 2 | 7.31 | |
| Superior temporal gyrus | R | 50 | -42 | 14 | 4.90 | |
| **Cluster 2** (*k* = 1158) | | | | | | < .001 |
| Precuneus | L | -6 | -50 | 58 | 5.15 | |
| Precuneus | R | 6 | -48 | 48 | 5.00 | |
| Precuneus | L | -12 | -68 | 60 | 4.40 | |
| **Cluster 3** (*k* = 973) | | | | | | < .001 |
| Middle occipital gyrus | L | -44 | -72 | 0 | 4.87 | |
| Middle occipital gyrus | L | -40 | -62 | 4 | 4.62 | |
| Middle occipital gyrus | L | -22 | -90 | 4 | 4.62 | |

*Note.* Significant clusters including hemisphere h, cluster size *k*, MNI coordinates x, y, z, *t* value and *p* value (whole brain, FWE-corrected at $p < .05$, cluster level, $k > 279$). Anatomical regions were labelled with SPM12's Anatomy toolbox (version 2.2c; Eickhoff et al., 2005)

Table S3

*fMRI whole brain results for the contrast other passive (incongruent > congruent).*

| Clusters and brain regions | h | *x* | *y* | *z* | *t* value | *p* value |
|---|---|---|---|---|---|---|
| **Cluster 1** (*k* = 795) | | | | | | < .001 |
| Middle occipital gyrus | L | -48 | -74 | 2 | 5.29 | |
| Superior temporal gyrus | L | -54 | -44 | 12 | 3.89 | |
| Middle temporal gyrus | L | -48 | -52 | 6 | 3.53 | |
| **Cluster 2** (*k* = 797) | | | | | | < .001 |
| Middle temporal gyrus | R | 50 | -64 | 2 | 5.27 | |

*Note.* Significant clusters including hemisphere h, cluster size *k*, MNI coordinates x, y, z, *t* value and *p* value (whole brain, FWE-corrected at $p < .05$, cluster level, $k > 279$). Anatomical regions were labelled with SPM12's Anatomy toolbox (version 2.2c; Eickhoff et al., 2005)

*Region of interest results*. Table S7 shows the ANOVA table of the region of interest data in the fMRI study.

Table S7

*ANOVA using extracted activation of two ROIs in rSMG and rTPJ.*

| Effects | *dfs* | *F* | $p_{(two\text{-}tailed)}$ | gen. $\eta^2$ |
|---|---|---|---|---|
| Group | 1, 40 | 1.17 | .286 | 0.009 |
| Congruence | 1, 40 | 16.89 | .002 | 0.005 |
| Target | 2, 80 | 2.26 | .111 | 0.018 |
| ROI | 1, 40 | 29.07 | < .001 | 0.113 |
| Group × Congruence | 1, 40 | 2.32 | .136 | < 0.001 |
| Group × Target | 2, 80 | 1.42 | .248 | 0.012 |
| Group × ROI | 1, 40 | 0.60 | .441 | 0.003 |
| Congruence × Target | 2, 80 | 2.99 | .056 | 0.002 |
| Congruence × ROI | 1, 40 | 23.75 | < .001 | 0.002 |
| Target × ROI | 2, 80 | 1.11 | .335 | 0.003 |
| Group × Congruence × Target | 2, 80 | 1.36 | .262 | < 0.001 |
| Group × Congruence × ROI | 1, 40 | 0.39 | .534 | < 0.001 |
| Group × Target × ROI | 2, 80 | 0.18 | .838 | < 0.001 |
| Congruence × Target × ROI | 2, 80 | 0.52 | .596 | < 0.001 |
| Group × Congruence × Target × ROI | 2, 80 | 5.45 | .006 | 0.001 |

*Note.* ROI = region of interest; rSMG = right supramarginal gyrus; rTPJ = right temporoparietal junction.
